# Reverse-zoonoses of 2009 H1N1 pandemic influenza A viruses and evolution in United States swine results in viruses with zoonotic potential

**DOI:** 10.1101/2022.12.15.520479

**Authors:** Alexey Markin, Giovana Ciacci Zanella, Zebulun W. Arendsee, Jianqiang Zhang, Karen M. Krueger, Phillip C. Gauger, Amy L. Vincent Baker, Tavis K. Anderson

**Affiliations:** Virus and Prion Research Unit, National Animal Disease Center, United States Department of Agriculture, Agricultural Research Service, Ames, IA, USA; Department of Veterinary Microbiology and Preventive Medicine, College of Veterinary Medicine, Iowa State University, Ames, IA, USA; Department of Veterinary Diagnostic and Production Animal Medicine, College of Veterinary Medicine, Iowa State University, Ames, IA, USA

**Keywords:** influenza A, swine, zoonoses, pandemic preparedness

## Abstract

The 2009 H1N1 pandemic (pdm09) lineage of influenza A virus (IAV) crosses interspecies barriers with frequent human-to-swine spillovers each year. These spillovers reassort and drift within swine populations, leading to genetically and antigenically novel IAV that represent a zoonotic threat. We quantified interspecies transmission of the pdm09 lineage, persistence in swine, and identified how evolution in swine impacted zoonotic risk. Human and swine pdm09 case counts between 2010 and 2020 were correlated and human pdm09 burden and circulation directly impacted the detection of pdm09 in pigs. However, there was a relative absence of pdm09 circulation in humans during the 2020-21 season that was not reflected in swine. During the 2020-21 season, most swine pdm09 detections originated from human-to-swine spillovers from the 2018-19 and 2019-20 seasons that persisted in swine. We identified contemporary swine pdm09 representatives of each persistent spillover and quantified cross-reactivity between human seasonal H1 vaccine strains and the swine strains using a panel of monovalent ferret antisera in hemagglutination inhibition (HI) assays. The swine pdm09s had variable antigenic reactivity to vaccine antisera, but each swine pdm09 clade exhibited significant reduction in cross-reactivity to one or more of the human seasonal vaccine strains. Further supporting zoonotic risk, we showed phylogenetic evidence for 17 swine-to-human transmission events of pdm09 from 2010 to 2021, 11 of which were not previously classified as variants, with each of the zoonotic cases associated with persistent circulation of pdm09 in pigs. These data demonstrate that reverse-zoonoses and evolution of pdm09 in swine results in viruses that are capable of zoonotic transmission and represent a potential pandemic threat.

**Author Summary:** The diversity and evolution of influenza A virus (IAV) in pigs is linked to the emergence of IAV with pandemic potential. Human-to-swine transmission of the 2009 H1N1 pandemic (pdm09) IAV lineage repeatedly occurred across the past decade and has increased genetic diversity in pigs: sporadic swine-to-human cases are associated with these viruses. We measured the frequency of human-to-swine transmission of the H1N1 pandemic IAV lineage between 2009 and 2021 and determined how this affected the diversity of IAV in swine and zoonotic risk. We detected 371 separate human-to-swine spillovers, with the frequency of interspecies transmission increasing when the burden of IAV was highest in the human population. Most spillovers were single events without sustained transmission, but a small subset resulted in the emergence, persistence, and cocirculation of different pdm09 genetic clades in US pigs. Each of the pdm09 representative of different persistent spillovers was genetically and antigenically different from human seasonal vaccine strains. The persistence of pdm09 within pigs resulted in at least five recent swine-to-human transmission events. These data suggest that controlling IAV infection in humans working with swine can minimize spillover into pigs, reduce resulting genetic diversity of IAV in pigs, and proactively reduce the potential for swine-to-human transmission of IAV with pandemic potential.

## 1. Introduction

Human-to-swine transmission of influenza A viruses (IAV) increases genetic and antigenic diversity of swine IAV, confounding control efforts and impacting animal health^1^. There are more than 20 genetic clades of hemagglutinin (HA) genes within the H1N1, H1N2, and H3N2 subtypes of IAV circulating in US swine, and some or all the 8 virus genes are derived from an ancestral human IAV^1–3^. A consequence of the observed genetic diversity is frequent gene reassortment and rapid evolution in the surface proteins^4^ resulting in antigenically drifted viruses to which the human population may have little to no immunity^5–7^. This dynamic results in endemic swine IAV that have the potential to transmit to humans and represent a zoonotic threat (Figure 1). The quintessential example for the importance of the human-swine IAV interface is the 2009 H1N1 pandemic: this HA lineage established in swine in the early 1900s concurrent with the 1918 human IAV pandemic, evolved in the swine host for over ninety years, and subsequently caused the first IAV pandemic of the 21^st^ century^8^.

**Figure 1.**
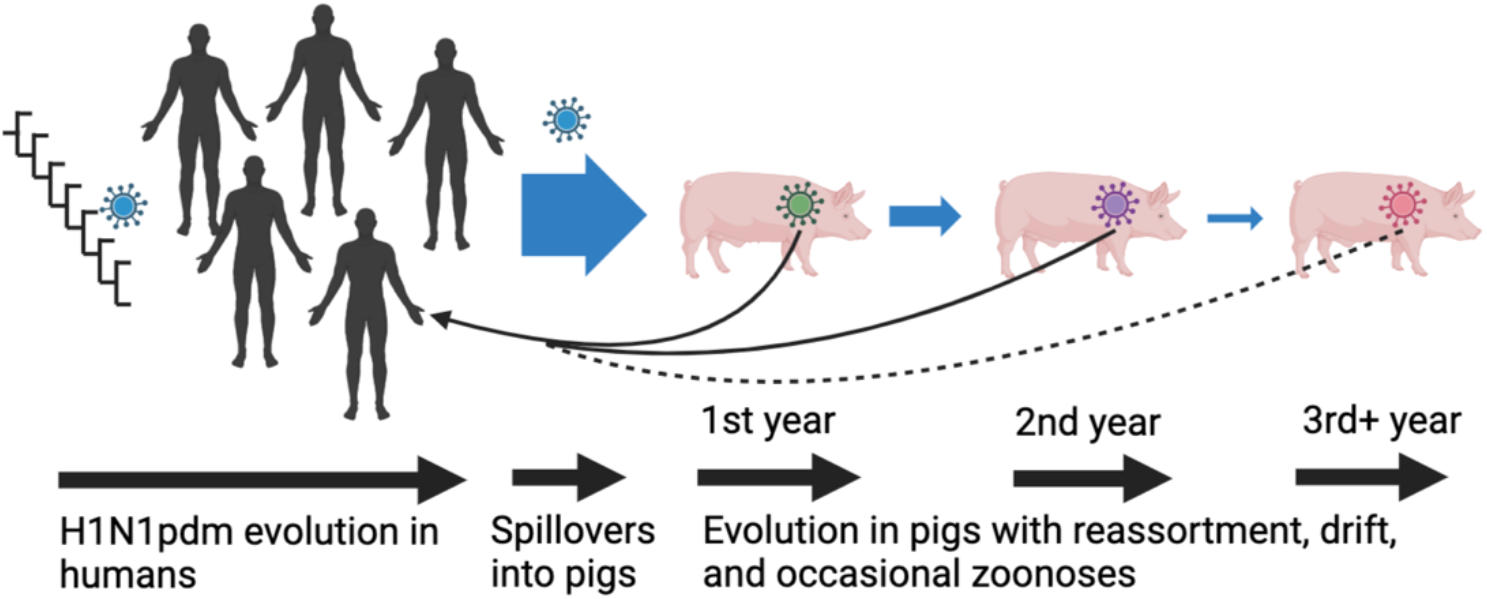
A conceptual model describing the human-swine interface and zoonotic risk for the 2009 H1N1 pandemic lineage (pdm09). There is a linear correlation between the burden of pdm09 in human populations and the number of reverse-zoonoses; a subset of these spillovers (indicated by smaller arrow width) persists for multiple years in pigs, evolve, and drift from the founding human-origin pdm09 strain; occasionally these persistent pdm09 cause zoonotic infections. These patterns were derived from pdm09 dynamics in the US.

The swine-origin 2009 H1N1 pandemic IAV (pdm09) replaced the previously circulating human seasonal H1 lineage^6,9^ and has continued to change the evolutionary landscape of IAV in both hosts. The pdm09 virus was a reassorted virus with gene segment origins from avian, human, and swine hosts; each of the gene segments had been evolving in swine for differing durations with data suggesting that the virus had been in pigs for ~5-10 years prior to emerging as a pandemic^9,10^. This multiple reassorted virus adapted to the human host and is now the human seasonal H1N1 with seasonal epidemic peaks and transmission^6^. There is also considerable support for this human IAV lineage acting as a major contributor to the genetic diversity of IAV in swine through reverse-zoonotic events^11–13^. Following a human-to-swine spillover, the pdm09 virus donates internal gene segments to endemic swine IAVs via reassortment^12–16^, and the surface proteins are maintained with pdm09 origin or they are paired with other endemic swine-origin internal genes^11,17^.

In this study we investigated interspecies transmission of the H1N1pdm09 lineage. We were motivated to understand whether the observed reduction in IAV circulation in the 2020-21 human flu season during the COVID-19 pandemic^18–20^ impacted the evolution of the pdm09 lineage in swine and how this affected its zoonotic potential. We applied a comprehensive phylogenetic analysis to human and swine pdm09 gene sequence data collected between 2009 and 2021 to show frequent human-to-swine spillovers of the H1N1pdm09 virus (approximately 370 independent events). The number of spillovers was significantly higher in seasons with high H1N1pdm09 circulation in the human population, and human-to-swine spillovers continuously fueled pdm09 circulation in swine. In the 2020-21 season with low human IAV circulation, the pdm09 remained a frequently detected HA genetic clade in US swine. We demonstrated that the 2020-21 pdm09 swine detections were the result of approximately 150 human-to-swine spillovers that occurred during the 18-19 and 19-20 flu seasons that subsequently circulated and evolved in swine populations. A significant proportion of these spillovers were sustained in US swine and continue to circulate and evolve. The contemporary swine pdm09 viruses showed a substantial reduction in cross-reactivity to one or more human H1N1pdm09 vaccine viruses. The adaptation and evolution of human IAV in swine may result in viruses with zoonotic potential, and our data revealed at least 17 zoonotic H1N1pdm09 transmission events between 2010 and 2021, 8 of which occurred in 2020 and 2021.

## 2. Methods

### Human and swine IAV detection frequency in the United States

The weekly human influenza detection report containing the number of positive samples each week in the United States was downloaded from WHO FluNet on September 9, 2021 (https://www.who.int/tools/flunet). The FluNet report displays the number of positive samples for the following IAV subtypes: A(H1N1)pdm09, other H1, H3, H5, unsubtyped. Between 2010 and 2021, there were no detections of H5 or non-A(H1N1)pdm09 H1 viruses; i.e. all IAV activity was due to A(H1N1)pdm09 and H3 subtypes. For a fixed season, to obtain an estimate for the A(H1N1)pdm09 detections in humans among unsubtyped IAV samples, we assumed that the A(H1N1)pdm09 to H3 ratio among the subtyped samples was representative of all influenza A detections, and split the number of unsubtyped samples according to that ratio. The swine IAV detection report was downloaded from Iowa State University FLUture on September 13, 2021^21^. The report contained daily detections for each (US) IAV in swine clade, including pdm09, beginning in 2009.

Using the FluNet and FLUture reports we aggregated the weekly and daily detection numbers into seasons, starting with the 2010-11 season. For human data, we used the standard Northern hemisphere season definition of October 1 to the following April 30^22^, and we considered all weeks that were fully contained in that interval for each season. For the respective swine season, we defined it as November 1 to the following October 31 since IAV circulates in swine year-round and used the respective FLUture records for aggregation. For example, the 2010-11 human season was defined as October 2010 to April 2011 and the swine season as November 2010 to October 2011. The start of the swine season was defined as November 1 to align the yearly detection peak of IAV in swine in September-October^23,24^ with the prior human season (Supplemental Figure S1). This seasonal IAV peak in swine precedes the seasonal IAV activity in humans, thus it is unlikely to be driven by same-season human-to-swine spillovers. Instead, the swine IAV detections in September-October are predominantly driven by continued circulation of pdm09 viruses through the summer months from prior seasonal human-to-swine spillovers.

### Collection of HA genetic sequence data for human and swine IAVs

Human hemagglutinin (HA) gene sequences of the H1N1 subtype collected between January 2009 and October 2021 were downloaded from GISAID EpiFlu database on March 14, 2022^25^. Swine HA gene sequences of the H1 subtype collected between January 2009 and October 2021 were downloaded from Influenza Research Database accessed on February 18, 2022^26^, and merged with previously unpublished sequences from the Iowa State University Veterinary Diagnostic Laboratory (accessions provided in Supplemental Table S1). We classified all collected H1 sequences using octoFLU^27^ and maintained only pdm09-like HAs. Sequences that did not have a month of collection or those less than 1690 nucleotides long were removed. This process resulted in 12,823 human HA genes and 1,112 swine HA gene sequences. We aligned all sequences with MAFFT v.7.222^28^, trimmed them to the coding region, and computed the number of substitutions (Hamming distance) between each pair of aligned human and swine sequences.

### Inferring interspecies transmission with phylogenetic analysis

We inferred a maximum likelihood phylogenetic tree with IQ-TREE v1.6.12^29^ under the GTR site substitution model with empirically estimated base frequencies and 5 free-rate categories^30,31^. We also generated a time-scaled phylogeny, rooted under the strict molecular clock assumption using TreeTime v0.8.4 phylodynamic toolkit^32^. The host state was inferred for each ancestral node (i.e., human/swine) on the time-scaled phylogeny using a TreeTime mugration model. Finally, we inferred ancestral amino acid substitutions in the HA1 protein domain on the rooted maximum likelihood tree with TreeTime. HA1 amino acid sequences for the tips were extracted from the HA genetic sequences using Flutile v.0.13.0 (https://github.com/flu-crew/flutile). We repeated the process using the same alignment 20 times to obtain confidence intervals associated with estimates of interspecies transmission in the downstream analyses. For all the statistics computed over the time-scaled phylogenies (as described below) we computed and report the median value and minimum-maximum ranges across 20 replicates.

### Identifying the seasonal origin of human-to-swine pdm09 spillovers and persistence of spillovers in US swine

To identify whether a swine pdm09 HA gene was a same-season or prior season spillover, we established the following process. We refer to a swine HA sequence from season *S* as a *same-season spillover*, if there was a human-to-swine spillover in the same season *S* and the swine sequence under consideration came from the transmission chain caused by that spillover. First, we used an inferred phylogenetic tree with host annotations on ancestral nodes to identify same-season spillovers among sequences from each season between 2010-11 and 2019-20. That is, we classified a swine sequence from season *S* as a same-season spillover if it is nested within a human clade with human sequences strictly from season *S* or later. We called a clade ‘human’ if the inferred host of its most recent common ancestor (MRCA) was a human host.

For the 2020-21 season, there were insufficient HA sequences isolated from humans in the US to apply the same phylogenetic method. Therefore, we applied rule mining to identify strong associations in the data prior to the 2020-21 season and used them to estimate the fraction of same-season spillovers in 2020-21. For rule mining, we considered all swine HA sequences from seasons 2013-14, 2014-15, 2015-16, 2016-17, 2017-18, 2018-19, and 2019-20 (a total of 705 sequences). We excluded seasons prior to 2013-14 due to the lower availability of swine pdm09 sequences. For each swine sequence collected in season *S* we then defined two predictor variables: (i) maximum similarity of that sequence to human sequences in season *S*-1, and (ii) maximum similarity to swine sequences in season *S*-1. For example, for a swine sequence from season 2018-19, we consider its similarity to human and swine sequences from the 2017-18 season. We then observed that for all same-season spillovers the maximum similarity to the prior human season was at least as large as the maximum similarity to the prior swine season.

For each human-to-swine spillover, i.e., a human-to-swine host transition along the time-scaled phylogeny, we inferred the season of that spillover and the duration that spillover persisted in swine in years. Each spillover was associated with a unique tree edge, where the parent had a human host and the child had a swine host. Then the date of spillover was assumed to be the estimated date of the parent node. We account for uncertainty in the spillover date estimation by including the few months prior to October and after April for the human season estimation. Specifically, if the spillover date was in or after July of year *Y*, then we inferred the spillover season as *Y*-(*Y*+1); otherwise, the spillover season was (*Y*-1)-*Y*. The length of the spillover was estimated as the height of the subtree rooted at the child node plus the length of the edge on a time-scaled phylogeny (Table 1).

**Table 1.**
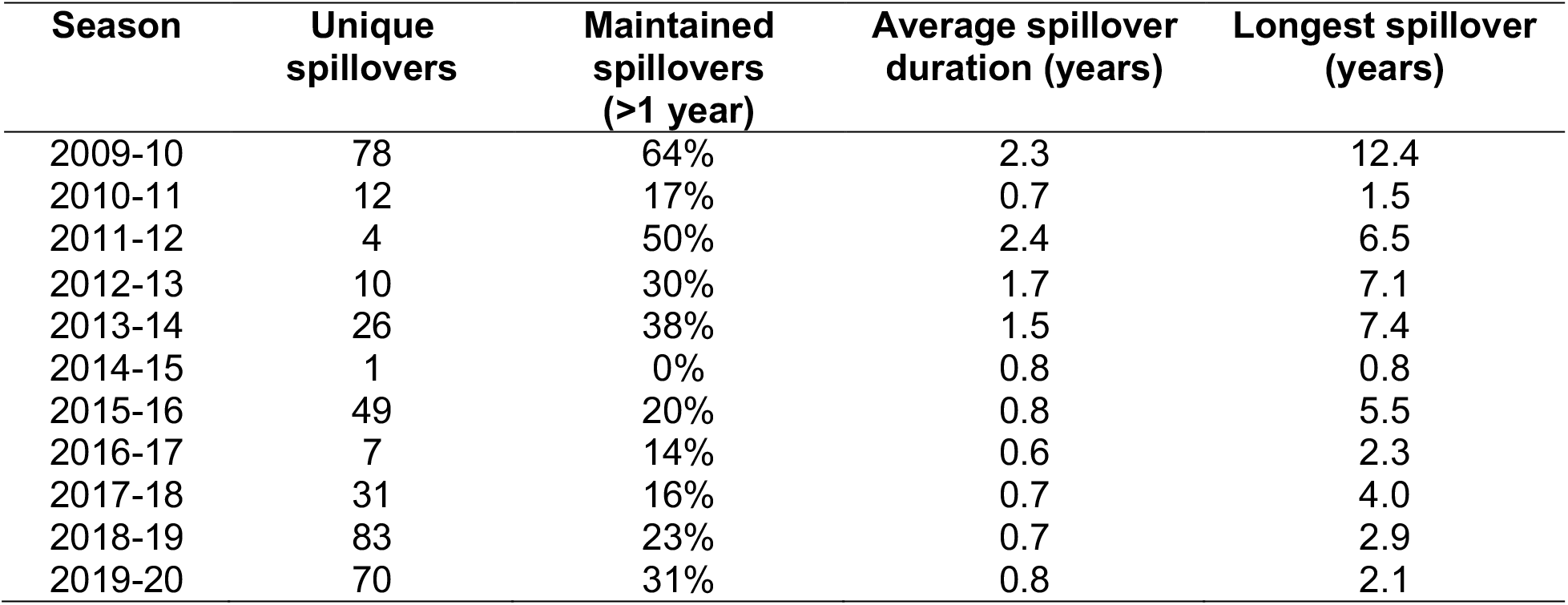
The number of human-to-swine spillovers grouped by season. Unique spillovers were identified using phylogenetic methods with the duration of spillover assessed using a temporally scaled phylogenetic tree. The proportion of spillovers that were maintained for longer than a year are indicated, along with the average and longest duration of detection.

| Season | Unique spillovers | Maintained spillovers (>1 year) | Average spillover duration (years) | Longest spillover (years) |
| --- | --- | --- | --- | --- |
| 2009-10 | 78 | 64% | 2.3 | 12.4 |
| 2010-11 | 12 | 17% | 0.7 | 1.5 |
| 2011-12 | 4 | 50% | 2.4 | 6.5 |
| 2012-13 | 10 | 30% | 1.7 | 7.1 |
| 2013-14 | 26 | 38% | 1.5 | 7.4 |
| 2014-15 | 1 | 0% | 0.8 | 0.8 |
| 2015-16 | 49 | 20% | 0.8 | 5.5 |
| 2016-17 | 7 | 14% | 0.6 | 2.3 |
| 2017-18 | 31 | 16% | 0.7 | 4.0 |
| 2018-19 | 83 | 23% | 0.7 | 2.9 |
| 2019-20 | 70 | 31% | 0.8 | 2.1 |

### Identifying putative swine-to-human transmission in 2020-21

Our phylogenetic analysis of interspecies transmission using the HA gene identified 16 putative variant infection cases resulting from swine-to-human transmission events in the pdm09 lineage between 2010 and October 2021. We considered a swine-to-human transmission event resulting in a putative variant infection if it had support of 20/20 across the 20 phylogenetic replicates. Of the 16 variant cases, 5 cases were confirmed and reported by the US Centers for Disease Control and Prevention (CDC) (https://gis.cdc.gov/grasp/fluview/Novel_Influenza.html), and these 5 had reassorted genomes with swine-endemic internal genes, but the remaining 11 were classified as human seasonal pdm09. There was an additional CDC-confirmed variant case with a reassorted genome (A/Iowa/02/2021, EPI_ISL_2479995): this strain was not detected in our HA phylogenetic analysis, as there were no similar swine HA genes, and the determination was based upon the reassorted genome. Consequently – there were a total of 17 putative or confirmed variant cases between 2010 and October 2021. In the 2020-21 human flu season, the putative (non-confirmed) variants were A/Iowa/22/2020 (EPI_ISL_613417), A/Iowa/23/2020 (EPI_ISL_766961), A/North Carolina/01/2021 (EPI_ISL_2519774), and A/Iowa/01/2021 (EPI_ISL_1459510). To support the assertion that these were swine-origin zoonotic cases, we downloaded all globally available human gene segments associated with the H1N1 subtype collected from 2019 to 2021 from GISAID^25^ – a total of 108,693 sequences [accessed December 8, 2022]. For swine, we downloaded gene segments collected in the US from 2019 to 2021 from the Influenza Research Database^26^ (8,808 sequences associated with H1 HA) [accessed December 8, 2022]. A BLAST^33^ database was built with all the collected sequences using NCBI BLAST+^34^. We used blastn (megablast with default gap scoring penalties) to identify genes that most closely matched A/Iowa/22/2020, A/Iowa/23/2020, A/North Carolina/01/2021, and A/lowa/01/2021 gene segments. For all 8 genes from each of the 4 strains, swine sequences were always among the top blast hits. For each of the 4 strains, at least 5 out of 8 genes were *strictly* closer to respective swine-origin genes than to human-origin genes, including 4 largest genes: PB2, PB1, PA, and HA. Only in two cases (MP genes from A/North Carolina/01/2021 and A/lowa/01/2021), there was a human gene sequence that was closer to the query gene than the closest swine gene sequence (1 substitution difference).

### Human vaccine virus reactivity to swine H1 derived from seasonal human-to-swine spillovers

Representative swine pdm09 strains reflecting distinct human-to-swine seasonal spillovers with sustained transmission in swine were identified. We extracted swine H1pdm sequences detected between July 2020 and October 2021 (n=251). We grouped these sequences by season of introduction to the swine population determined in the phylogenetic analysis above. Within each group we computed a consensus HA1 amino acid sequence and identified the swine strain within the USDA IAV in swine virus repository with the highest identity to the consensus. This process resulted in 5 strains that came from distinct spillovers in seasons 2013-14, 2015-16, 2017-18, 2018-19, and 2019-20 (see Supplementary Table 1). Due to cocirculation of at least two clades of swine pdm09 from the 2019-20 season, we selected an additional strain: this clade included the putative variant cases A/North Carolina/01/2021, and A/Iowa/01/2021. The 6 strains were tested against a panel of ferret antisera generated to human H1 vaccines. Ferret sera to human pdm09 seasonal vaccine strains (A/California/04/2009, A/Michigan/45/2015, A/Brisbane/2/2018, A/Wisconsin/588/2019) were provided by the Virology, Surveillance and Diagnosis Branch, Influenza Division, Centers for Disease Control and Prevention (CDC), Atlanta, Georgia, or generated at the National Animal Disease Center, USDA-ARS (A/Hawaii/70/2019). Ferrets were cared for in compliance with the Institutional Animal Care and Use Committee of the National Animal Disease Center, USDA-ARS. Ferret antisera were heat inactivated at 56°C for 30 min, then treated with a 20% suspension of kaolin (Sigma–Aldrich, St. Louis, MO) followed by adsorption with 0.75% guinea pig red blood cells (RBCs). HI assays were performed according to standard techniques. When available, two biological replicates of each virus anti-sera were used. Geometric mean titers obtained by log2 transformation of reciprocal titers were used for the comparison.

## 3. Results

### Human-to-swine transmission of the H1N1 pandemic lineage occurs frequently

We inferred the ancestral host state for 13,935 human and swine pdm09 HA genes sequenced in the US between 2009 and 2021 (Figure 2)^32^. This analysis estimated that pdm09 viruses were introduced from humans into pigs around 371 times (369-380 range) in the US following the emergence and establishment of the pdm09 as the H1 human seasonal lineage. In reverse, swine-to-human transmission resulted in 17 zoonotic pdm09 cases since 2010. 11 of these putative zoonotic cases, despite phylogenetic evidence, were not previously identified, likely because the sequenced genomes had high nucleotide similarity to other human-seasonal pdm09 viruses (at least 99.2% identity across all genes). Notably, some pdm09 clades that were replaced and disappeared in the human population continued to circulate and evolve in swine (e.g., clades 6b1.A1 and 6b1.A7) evidenced by swine IAV detected in 2021 but seeded from earlier human pdm09 seasons. Of the pdm09 introduced into swine, ~45% were maintained longer than a year (Table 1), with a 1-year average duration of circulation following spillover. Of the pdm09 in swine with whole genome sequence data (n=211), more than 50% maintained pdm09 lineage internal genes, but 49% of the strains had reassorted genomes, with the proportion of internal gene patterns mirroring the primary patterns detected across all H1N1 and H1N2 sequenced strains (Figure S2).

**Figure 2.**
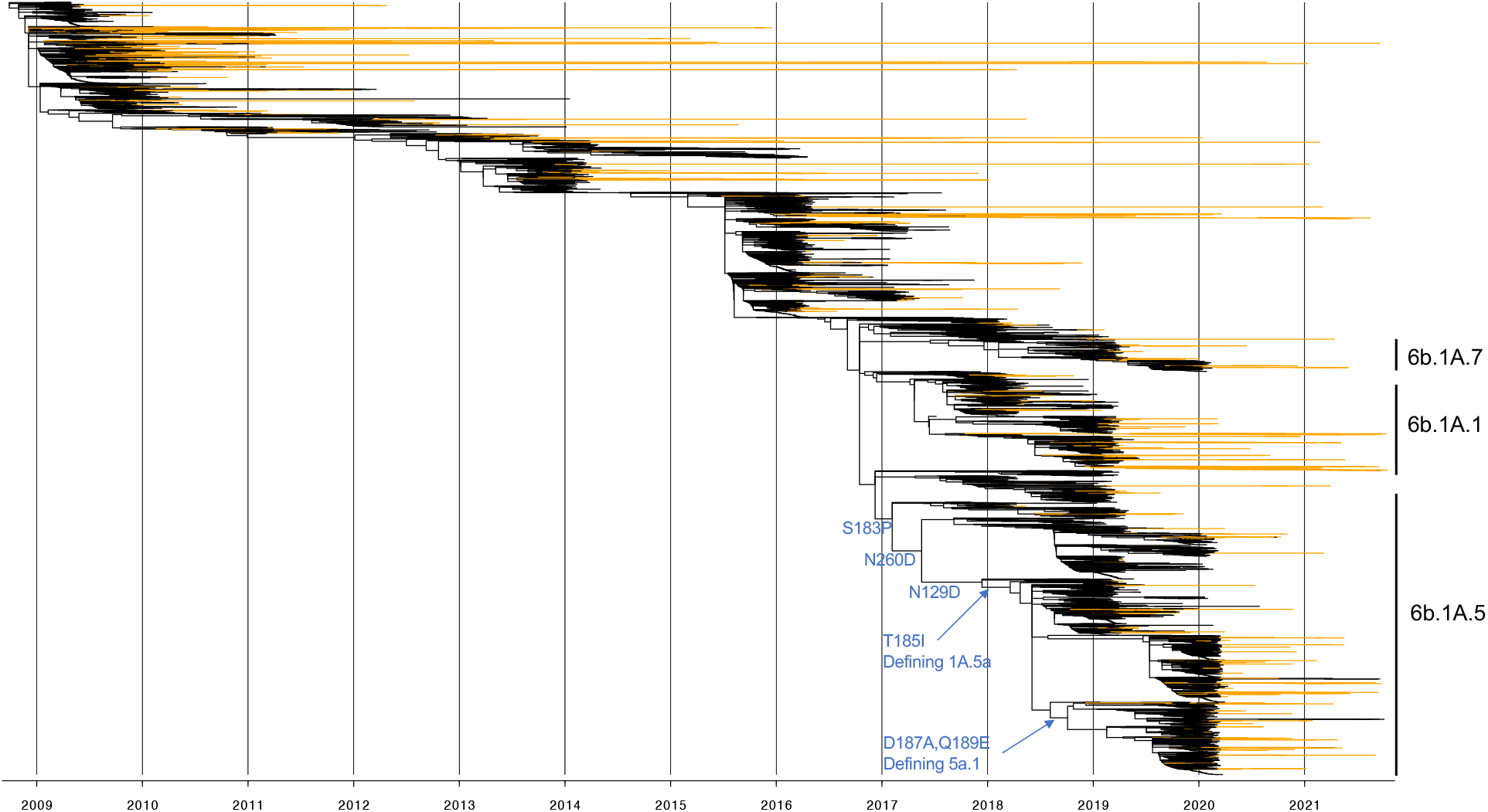
Time-scaled phylogeny of 13,935 swine and human hemagglutinin (HA) genes within the 2009 H1N1 pandemic lineage (pdm09) collected between 2009 and 2021. Orange branches indicate evolution in swine and black branches in humans. Human H1N1pdm09 seasonal clade names are annotated and clade defining amino acid mutations and those associated with antigenic change are annotated across the trunk of the phylogeny.

### Seasonal fluctuations in human pdm09 IAV burden determined the frequency of detection of swine pdm09 between 2010 to 2020

The aggregated pdm09 detection statistics for human and for swine data demonstrated a strong positive linear correlation for the 10 flu seasons 2010-11 to 2019-20 (Figure 3A, linear regression, *R*^2^ = 0.9; Pearson correlation coefficient = 0.95). Consequently, it was possible to estimate the number of pdm09 HA genes detected in swine in any given year using the human pdm09 burden. Consistent with prior reporting^11^, the observed number of human-to-swine spillovers was an order of magnitude higher than the number of swine-to-human spillovers. Therefore, the observed correlation between detection numbers was largely driven by human-to-swine spillovers, and these spillovers were detected more frequently in seasons with higher pdm09 burden in humans (Figure 3B).

**Figure 3.**
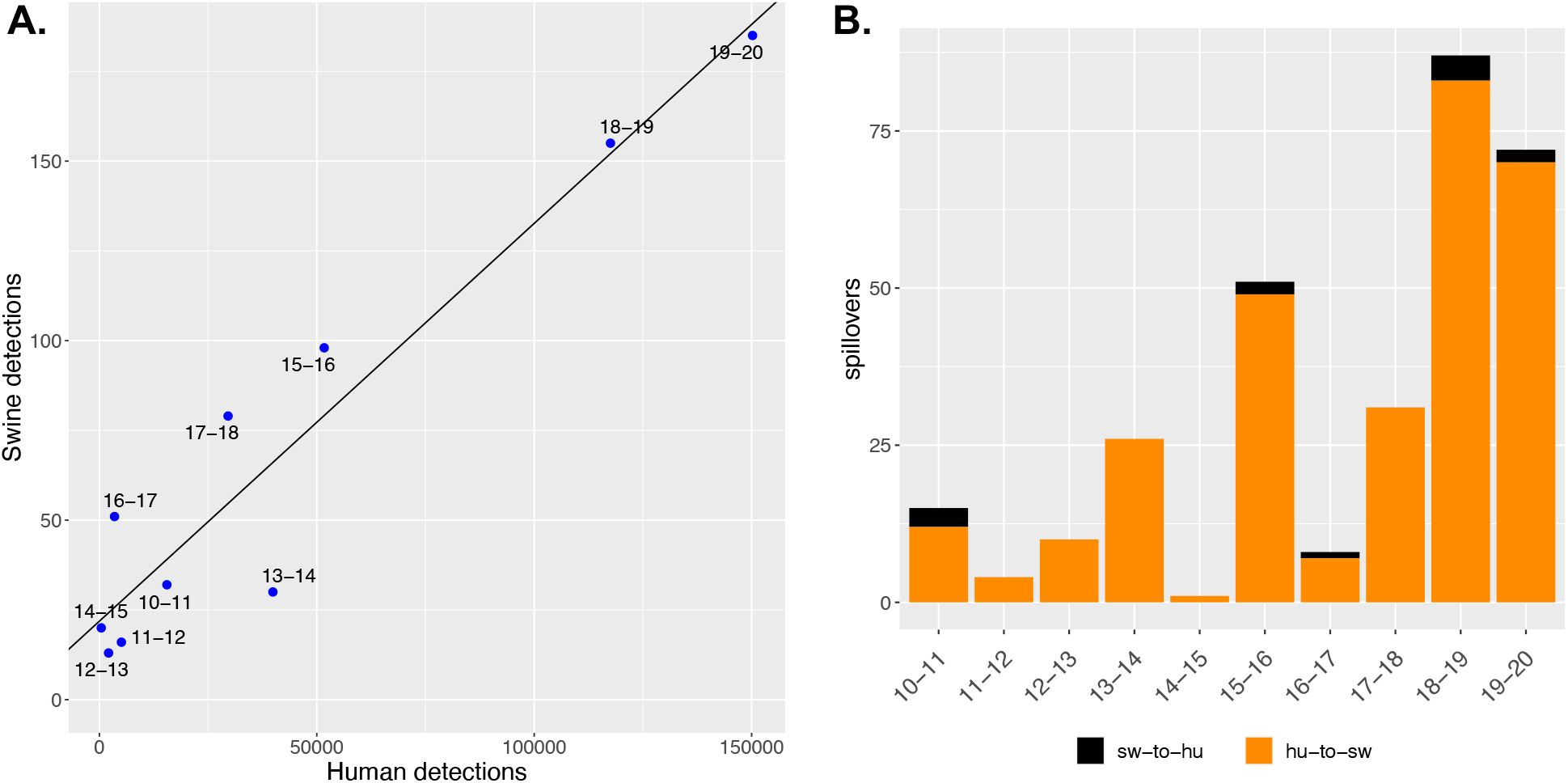
Association between human and swine H1pdm09 detection frequency. (A) H1N1pdm09 detections in humans and swine for each season between 2010-11 and 2019-20. The linear regression line was fitted with ordinary least squares and each blue dot is labeled by the corresponding influenza season. The 2020-21 season was omitted due to low detection of H1N1pdm09 in human populations. (B) The number of interspecies transmission events grouped by influenza season and host origin, human-to-swine (hu-to-sw) in orange and swine-to-human (sw-to-hu) in black.

The phylogenetic evidence for interspecies transmission was complemented by analyzing the temporal relationship between human and swine pdm09 detections. We analyzed swine and human pdm09 HA sequences collected in seasons between 2011-12 and 2018-19 (excluding the ‘border’ 2010-11 and 2019-20 seasons to ensure high-confidence data for the prior and future seasons). For each swine HA sequence, we determined the most genetically similar human HA sequence and compared the collection date for human and swine samples. If there were multiple human HA sequences with the same genetic distance to the swine sequence, we selected the sequence with the latest collection date. For 86% of the swine HA gene sequences, the closest human HA sequence was collected prior to the swine sequence. These data support the proposition that the primary direction of interspecies transmission of the pdm09 lineage was human-to-swine. Similarly, the rise in detections of pdm09 in humans consistently pre-dated the respective rise of pdm09 detections in swine across each of the seasons between 2010 and 2021 (Supplemental Figure S1).

### Detection of swine pdm09 in 2020-21 was not associated with new human-to-swine spillovers

The single season that did not follow the linear correlation between human and swine pdm09 detections was 2020-21. This season had record low human influenza activity in the US (18 confirmed pdm09 cases), and the regression line from Figure 3A would suggest very few swine pdm09 detections that season. However, swine pdm09 detections were high in 2020-21, exceeding the 2018-19 season, with 183 detections. We could not apply phylogenetic methods to examine potential human-to-swine spillovers in 2020-21 due to the low number of human pdm09 sequences from that season. However, using information from prior seasons we observed the following strong association: if a swine HA sequence was collected in season *S* and it came from a human-to-swine spillover that also occurred in season *S*, then it is more similar to the human sequences in season *S*-1 than to the swine sequences in season *S*-1 (Figure 4). These data show that there were only few (if any) human-to-swine spillovers in the 2020-21 season. Most of the available swine sequences (at least 158/175) from the 2019-20 season were the result of transmission of pdm09 in swine that originated as human-to-swine spillovers in 2019-20 or earlier. Our phylogenetic analysis showed that 46% (81/175) of these viruses originated from 2019-20 spillovers; the 2018-19, 2017-18, and 2015-16 seasons contributed 31%, 10%, and 11% of the detections, respectively. The five remaining detections were the result of sustained circulation of pdm09 in pigs from spillovers in the 2013-14 or 2009-10 seasons. Some spillovers established in the pig population with sustained transmission for more than 10 years (Figure 2; Table 1). Spillovers with over 5 years of circulation in US swine were detected across three or more US states.

**Figure 4.**
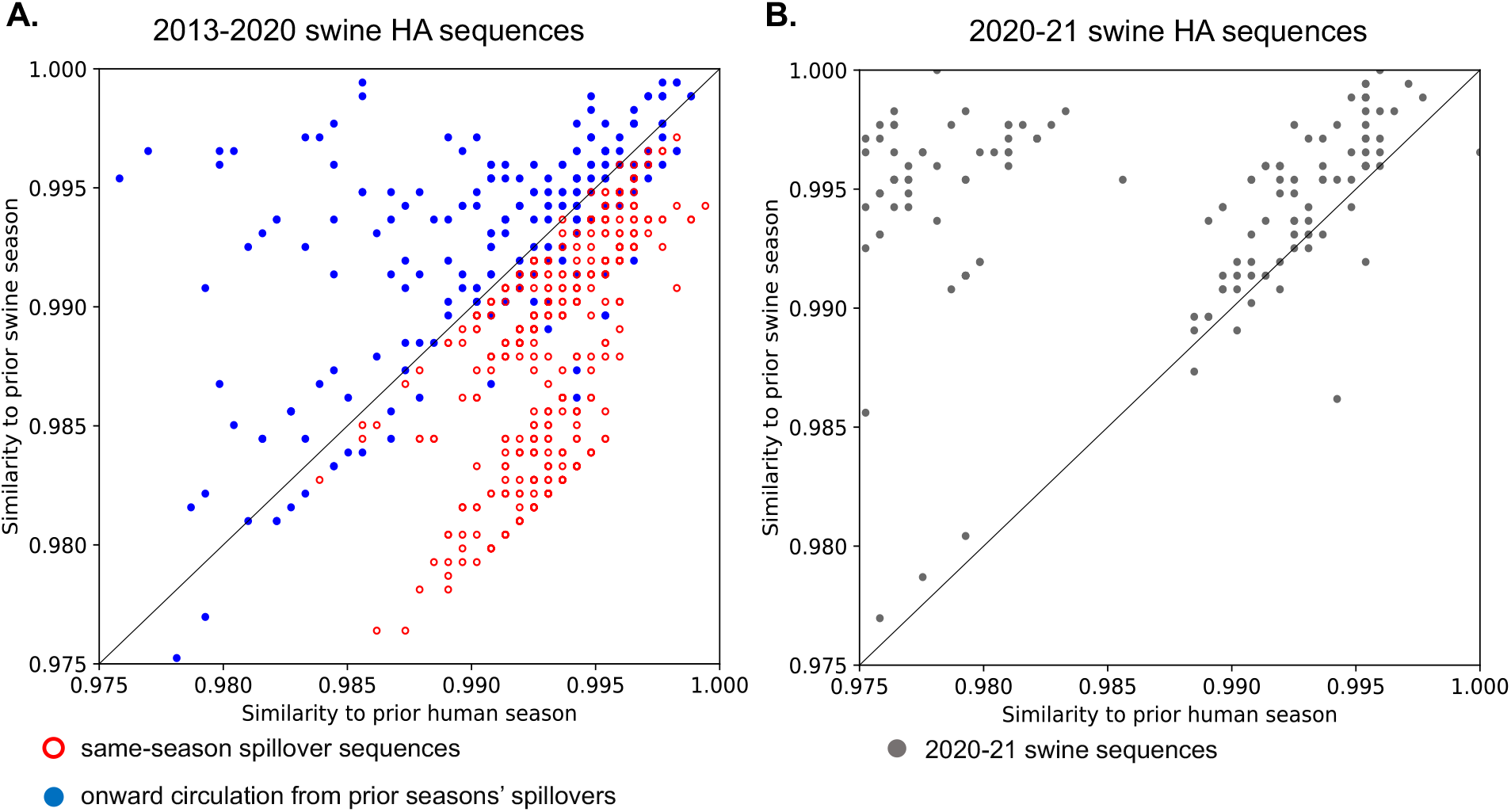
Rule mining to determine the host origin of H1N1pdm09 in swine measured in similarity in the hemagglutinin (HA) gene. (A) Swine HA sequences collected between the 2013-14 and 2019-20 seasons (inclusively) were colored red (open circle) if it was collected in season S and was derived from a human-to-swine spillover that happened in same season S. These data were colored blue (solid circle) if they reflected onward transmission of endemic swine pdm09 clades. The x-axis indicates the maximum similarity to a human HA sequence collected in season S-1 and the y-axis reflects maximum similarity to a swine HA sequence from season S-1. All same-season human-to-swine transmission events (open red circles) are located below the diagonal. (B) The 2020-21 season swine HA sequences plotted using the same genetic similarity thresholds with most data located above the diagonal reflecting onward transmission of endemic pdm09 in swine.

### Increased circulation of pdm09 in swine was associated with zoonotic detections

We investigated evidence for swine-origin pdm09 cases in humans (variant IAV) following the decrease in human seasonal IAV transmission in 2020-21 coincident with the COVID-19 pandemic. We identified 8 putative cases of swine-to-human transmission between September 2020 and October 2021, which represented almost half of all variant cases since 2010. In the 2020-21 season with relative absence of human seasonal IAV circulation, there were 5 publicly available genomes of pdm09 IAVs isolated from humans in the US between September 2020 and April 2021. One strain, A/Iowa/02/2021 (EPI_ISL_2479995), was a CDC-confirmed swine-origin infection with a reassorted pdm09 virus (with acquired triple-reassorted PB2 and NS genes; PB1 gene was not available). The other 4 strains had pdm09-lineage gene segments, and we provide phylogenetic evidence for zoonoses (Figure 5). On our phylogeny of 13,935 HA genes with inferred ancestral host state, all 4 HA genes were nested within swine pdm09 clades (Figure 5). The A/Iowa/22/2020 (EPI_ISL_613417) and A/Iowa/23/2020 (EPI_ISL_766961) strains were 99.95% identical to swine isolate A/swine/Iowa/A02524671/2020 (EPI_ISL_697636) across the entire genome with at most 2 nucleotide differences in individual genes and shared common amino acid mutations in HA1 region, including a mutation K142R in the Ca_2_ epitope (Figure 5B). The three strains A/Iowa/22/2020, A/Iowa/23/2020, and A/swine/Iowa/A02524671/2020 were collected within an 8-day interval in September 2020.

**Figure 5.**
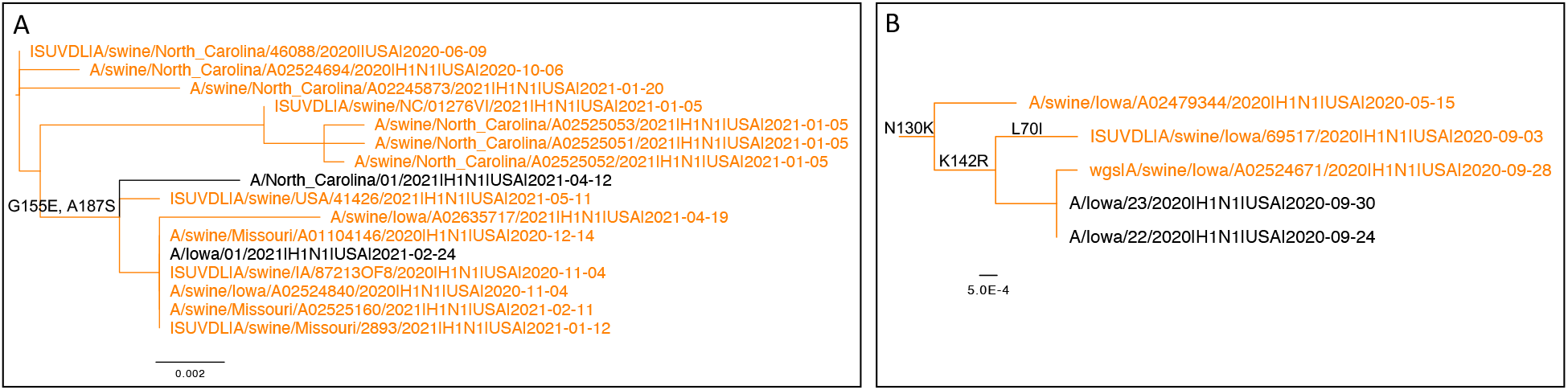
Subtrees extracted from an H1N1pdm09 phylogeny of 13,935 hemagglutinin (HA) genes demonstrating evidence for zoonoses. Swine HA genes are colored in orange and human HA genes are colored in black. Amino acid substitutions associated with phenotypic change are annotated on the tree edges. (A) Human HA genes, A/North Carolina/01/2021 and A/Iowa/01/2021, nested within a clade of 2019-20 and 2020-21 swine pdm09 HAs. (B) Human HA genes, A/Iowa/22/2020 and A/Iowa/23/2020, nested within a clade of 2019-20 swine pdm09 HAs. Edge lengths represent sequence divergence.

The A/North Carolina/01/2021 and A/Iowa/01/2021 HA were nested in a different swine pdm09 clade in the phylogeny (Figure 5A) and both strains were more closely related to respective swine-isolated genes than to the human-isolated genes. The two zoonotic strains had both acquired an HA1 mutation G155E (Figure 5A), which has been associated with binding to α2,6-linked glycans in combination with N129D mutation (a mutation shared by both isolates) and has been associated with antigenic escape^35,36^. The G155E mutation is present in 12% of the swine pdm09 sequences and only in 0.2% of human sequences in our dataset, it occurred independently 76 times throughout the evolution of the virus and is a repeatable amino acid change associated with the pdm09 HA gene in swine following human-to-swine transmission.

There were three additional swine-origin pdm09 variant infections in September 2021 that were confirmed by CDC: A/Iowa/05/2021 (EPI_ISL_7334894), A/Iowa/06/2021 (EPI_ISL_7334901) and A/North Dakota/12226/2021 (EPI_ISL_4702772). Our phylogenetic analysis suggests that these cases were a result of at least two swine-to-human transmissions within a single clade of swine IAVs that had been evolving in US swine since the beginning of 2019 and persisted (Figure S3). The HA of these three variant cases acquired 4 amino acid mutations in the HA1 globular head region, V47I, H51N, S85P, and R205K. Amino acid position 205 (in H1pdm numbering) is within the receptor binding site, the 190-helix^37^, and has been associated with antigenic escape^38^.

### Contemporary swine pdm09 strains exhibited wide variation in cross-reactivity with the human H1N1pdm09 vaccines

Representative contemporary swine pdm09 strains from different human-to-swine persistent clades were tested against reference ferret antisera generated against Northern Hemisphere human seasonal H1N1 vaccine strains (A/California/04/2009, A/Michigan/45/2015, A/Brisbane/2/2018, A/Wisconsin/588/2019, and A/Hawaii/70/2019) (Figure 6). The HA genes of the selected swine strains had circulated and evolved in US pigs for 1 to 7 years following different human-to-swine spillovers (Supplementary Table 1). All of the swine strains had HA and neuraminidase genes derived from the pdm09 lineage. The strains A/swine/Indiana/A02525081/2021 (2013-14 spillover) and A/swine/Missouri/A01104146/2020 (2019-20 spillover) maintained pdm09 lineage internal genes. The remaining 4 swine strains from the 2015-16, 2017-18, 2018-19, and 2019-20 contained reassorted internal genes derived from endemic swine lineages^39^ (Supplementary Tables 1-2). For each human pdm09 vaccine strain, there was one or more swine strains with at least 4-fold drop in cross-reactivity with the ferret antisera. Two contemporary swine strains, A/swine/Kansas/A02248038/2021 and A/swine/Missouri/A01104146/2020, ranged from 1.5 to 4 antigenic units (log2 titer ratio) away from all the vaccine antisera. Overall, the swine strains exhibited the largest drop in cross-reactivity with the A/Wisconsin/588/2019 vaccine strain, which was a WHO recommended vaccine strain for the 2021-22 and 2022-23 flu seasons.

**Figure 6.**
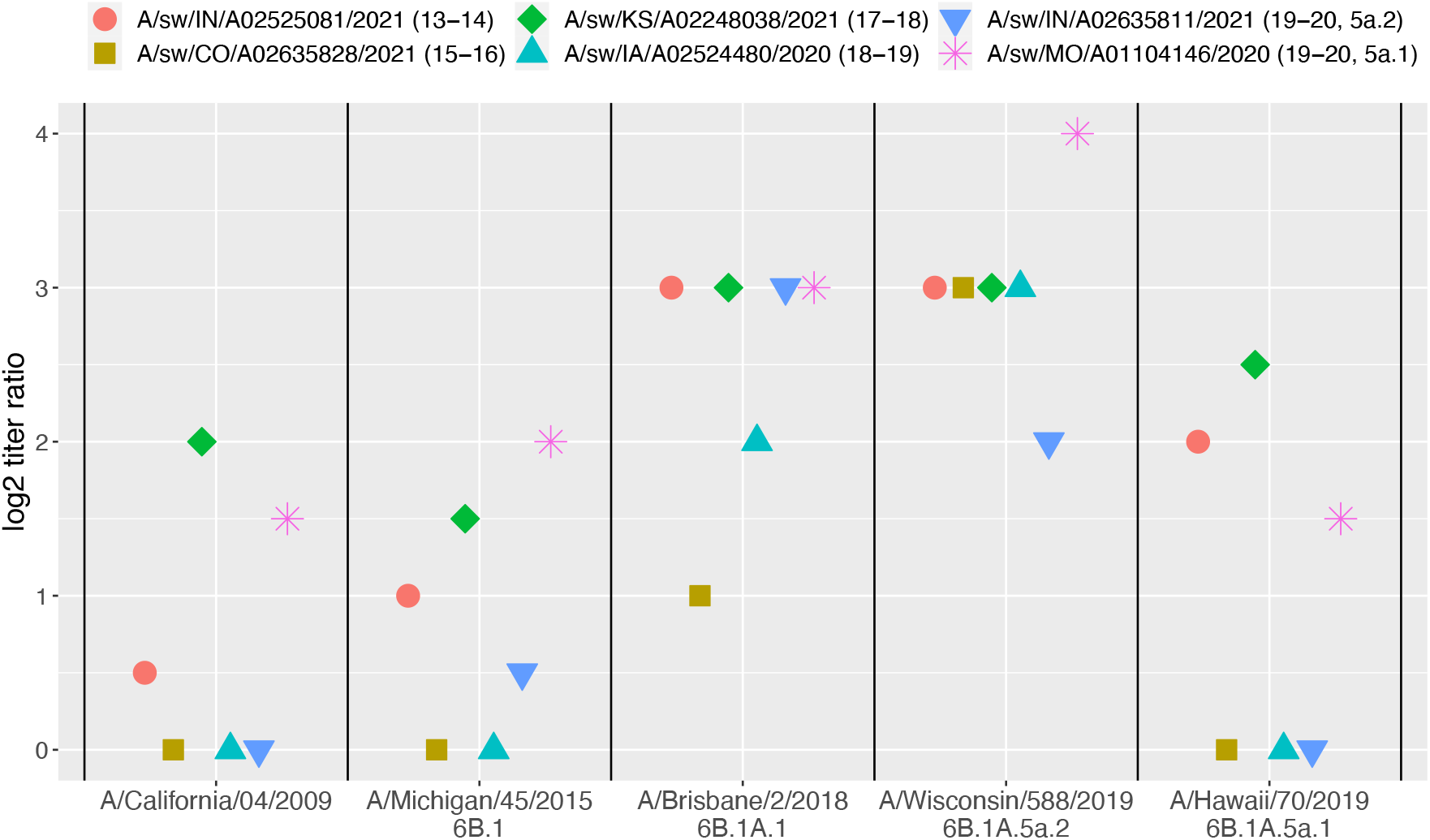
Hemagglutinin inhibition of contemporary swine pdm09 strains by pdm09 seasonal vaccine ferret antisera. The graph shows the antigenic distances as log2 of the ratio of homologous/heterologous geometric mean reciprocal titers. Three units of antigenic distance is equivalent to an 8-fold loss in HI cross-reactivity.

## 4. Discussion

Over the previous century, pandemic influenza A viruses (IAV) emerged through the transmission of IAV from animals to humans^40^; reassortment of IAV between human, avian, and swine influenza viruses^41^; or through recycled human viruses that circulated at a previous time^42^. We analyzed interspecies transmission patterns of the H1N1pdm09 (pdm09) IAV in the US since 2009. Our analysis confirmed that pdm09 frequently crosses the interspecies barrier between humans and swine. Following a human-to-swine spillover, a large fraction were maintained in US swine for longer than a year (~45%), more than 50% of these spillovers reassorted with endemic swine IAV^39^, and persistence and adaptation in the swine host resulted in viruses antigenically drifted from human seasonal vaccine strains. Our data demonstrate that pdm09 viruses established in US swine have drifted away from human seasonal vaccine strains, potentially reducing human population immunity to them. There is, consequently, a pressing need to characterize the breadth of genetic and antigenic diversity of pdm09 viruses circulating in swine to understand how transmission and evolution in this host impacts pandemic risk.

Our analysis demonstrated ~371 independent human-to-swine spillovers of the virus and 16 zoonotic infections after 2009. Most human-to-swine spillovers after 2010 (n=149) occurred in the 2018-19 and 2019-20 seasons, the seasons with the highest H1N1pdm09 burden in humans in the US. The number of human pdm09 detections more than doubled in the 2018-19 flu season when compared to the previous record-high season, 2015-16 (117,535 vs. 51,700 cases), and by another 28% in 19-20 (150,212 cases). The surge in pdm09 circulation led to substantially more human-to-swine spillovers and, consequently, increased pdm09 circulation in swine populations. This scenario reflects a “propagule-pressure” effect^43^. When there is a higher burden of pdm09 in the human population, it is more likely a human-to-swine spillover event occurs, and if there are more spillovers, one may have a genetic trait that facilitates establishment and transmission in the new host. This propagule effect is supported by an increased fraction of sustained human-to-swine spillovers in the 2018-19 and 2019-20 seasons. It is difficult to determine whether this dynamic affects zoonotic transmission given sparse but improved surveillance for IAV in swine and the role of the human-swine interface in variant detections^44^. However, we identified 8 variant infections in 2020-21 caused by pdm09 in swine originally derived from human-to-swine transmission during the prior 2019-20 season, and these variant cases occurred when the pdm09 clade was ~10% of all IAV in swine detections. In fact, all of the publicly available sequences from pdm09 human infections in the U.S. 2020-21 flu season were likely the result of swine-to-human spillovers. Four of those cases were infections with non-reassorted IAV. This suggests that swine can serve as a reservoir species for pdm09 viruses, swine-to-human spillovers may lead to re-appearance of older pdm09 clades in humans, and that pdm09 genetic clades frequently detected in commercial swine should be considered within zoonotic risk assessments.

In the 2020-21 season, there was high detection of pdm09 in US pigs, despite the absence of new human-to-swine spillovers that seeded detections in prior seasons. In general, only a fraction of human-to-swine spillovers were maintained into the next flu season and further. However, the 2018-19, and 2019-20 seasons exhibited an increase in the proportion of persistent pdm09 clades in swine. This dynamic could have occurred for three potential reasons: intrinsic virus factors associated with transmission; ecological factors associated with swine production; or the host immune profile. Though there are no consistent virological factors predictive of IAV transmission in swine, prior research has demonstrated that certain internal gene combinations are more frequently detected than others^45^, and certain gene segments appear to influence detection frequency^46^. Our data detected that 34% of pdm09 acquired internal genes via reassortment from endemic swine IAV, and 10% acquired a genome constellation associated with higher transmission when paired with an H3 subtype^47^. A second factor influencing pdm09 persistence are agricultural practices; populations of breeding, weaning, and growing swine are moved across the US and occasionally mingle and exchange viruses. These production practices facilitate the persistence of minor populations of IAV genes^48^ and as pigs move, the minor populations of genes are introduced to different regions and naïve host populations^49,50^ that affect subsequent detection patterns and reassortment^46^. The establishment and persistence of pdm09 in swine is also affected by the immunological profile of each population that is exposed to each virus. This profile is affected by pdm09 exposure in prior seasons, vaccination history that may or may not include a pdm09 component, and the number of distinct pdm09 lineage introductions to pigs in the same season. It is plausible that the absence of new pdm09 human-to-swine transmissions in 2020-21 may have resulted in a competitive release where waning population immunity to pdm09 provided more opportunity for prior season spillovers to persist.

A major implication of our research is that there was minimal amount of evolution in the HA of pdm09 viruses in swine following spillover, resulting in endemic pdm09 swine viruses that retain the potential to infect humans. Contrasting prior research^51^, the zoonotic cases we identified did not have a consistent swine signature, instead they generally maintained the genomic diversity within the HA gene of the founding human seasonal strains. The exception to this were the variants detected in 2020-21: these were associated with a G155E mutation that was detected in 12% of swine pdm09 genes, and that had arisen across the phylogeny at least 76 times. This amino acid was implicated as playing a major role in the antigenic variation of swine H1 viruses^36^. The repeated emergence and persistence of the G155E following human-to-swine transmission events is problematic as this single mutation may reduce the efficacy of human seasonal vaccines, and preexisting immunity to pdm09 may not prevent infection from a swine-origin pdm09. An additional 4 amino acid mutations in the HA1 globular head region, V47I, H51N, S85P, and R205K were also associated with variant cases in 2020-21. Of these, position 205 (in H1pdm numbering) is within the receptor binding site and predicted to be within an epitope^52^ and was associated with antigenic escape^38^. Similarly, position 47, 51, and 85 have been predicted to be associated with epitope E^52^, and the accumulated number of mutations in this epitope in swine pdm09 may impact antibody binding in human populations. In each of the variant cases, the founding human seasonal pdm09 event occurred in 2019, suggesting that within 2 years the viruses accrued mutations in critical HA1 regions than are likely to reduce the efficacy of current human vaccine immunity. This proposition is supported by our data that demonstrated the representative swine pdm09 from 2019-20 had a significant loss in reactivity to A/Brisbane/02/2018 (6B.1A.1) and A/Wisconsin/588/2019 (6B.1A.5a.2) vaccine strains, and one of the swine pdm09 had reduced reactivity to the A/Hawaii/70/2019 (6B1A.5a.1) vaccine. Generally, our HI data demonstrated a range of reactivities for each of the swine pdm09 to the five vaccine strains; this suggests that understanding pandemic risk requires characterization against a panel of vaccines as well as human serology data from different populations to understand how the immunological landscape in humans affects zoonotic risk^7,53^,

With each human-to-swine spillover of a pdm09, there is a new opportunity for the virus to antigenically drift and/or reassort with endemic swine viruses, creating a dynamic zoonotic risk profile. We demonstrated that the likelihood of a human-to-swine transmission event increases in seasons with higher pdm09 burden in humans, and the absence of widespread human pdm09 circulation in 2020-21 facilitated the persistence, spread, and evolution of pdm09 in swine. This resulted in endemic swine pdm09 lineages that accrued mutations in antigenic regions of the HA and human seasonal vaccines may lose efficacy against some of the pdm09 circulating in swine. Consequently, there is a need for continued genomic surveillance paired with characterization of IAV that capture the diversity of pdm09 in swine, as these maintain the potential for transmission to humans as they evolve. Ongoing surveillance is particularly relevant in light of the current highly pathogenic avian influenza (HPAI) H5N1 outbreak in North America^54^; if pigs could be infected with HPAI, reassortment with pdm09 in swine may result in viruses with increased host range and transmissibility in mammals^55^. This work demonstrates how a one health perspective is beneficial and that IAV pandemic planning requires the regular assessment of swine IAV to minimize the potential for another swine-origin pandemic.

## Data accessibility

All data, scripts, and the acknowledgment for GISAID sequences used in this manuscript are available at https://github.com/flu-crew/pdm-spillovers

## Authors’ contributions

AM: conceptualization, data curation, formal analysis, investigation, methodology, validation, visualization, writing-original draft, writing-review & editing; GCZ: formal analysis, writing-review & editing; ZWA: data curation, formal analysis; JZ: data curation, resources; KMK: data curation, resources; PCG: data curation, resources, funding acquisition, writing-review & editing; ALVB: conceptualization, funding acquisition, validation, writing-review & editing; TKA: conceptualization, funding acquisition, project administration, supervision, validation, writing-original draft, writing-review & editing. All authors gave final approval for publication and agreed to be held accountable for the work performed therein.

## Acknowledgements

We gratefully acknowledge pork producers, swine veterinarians, and laboratories for participating in the USDA Influenza A Virus in Swine Surveillance System and publicly sharing sequences. We thank C. Todd Davis and members of the Virology, Surveillance, and Diagnosis Branch, Influenza Division, Centers for Disease Control and Prevention, for provision of reagents used in these studies. We thank Scott Hensley from the University of Pennsylvania for the A/Hawaii/70/2019 virus. We also gratefully acknowledge all data contributors, i.e., the Authors and their Originating laboratories responsible for obtaining the specimens, and their Submitting laboratories for generating the genetic sequence and metadata and sharing via the GISAID Initiative, on which components of this research is based.

