## Supplemental Material for "Reverse-zoonoses of 2009 H1N1 pandemic influenza A viruses and evolution in United States swine results in viruses with zoonotic potential"

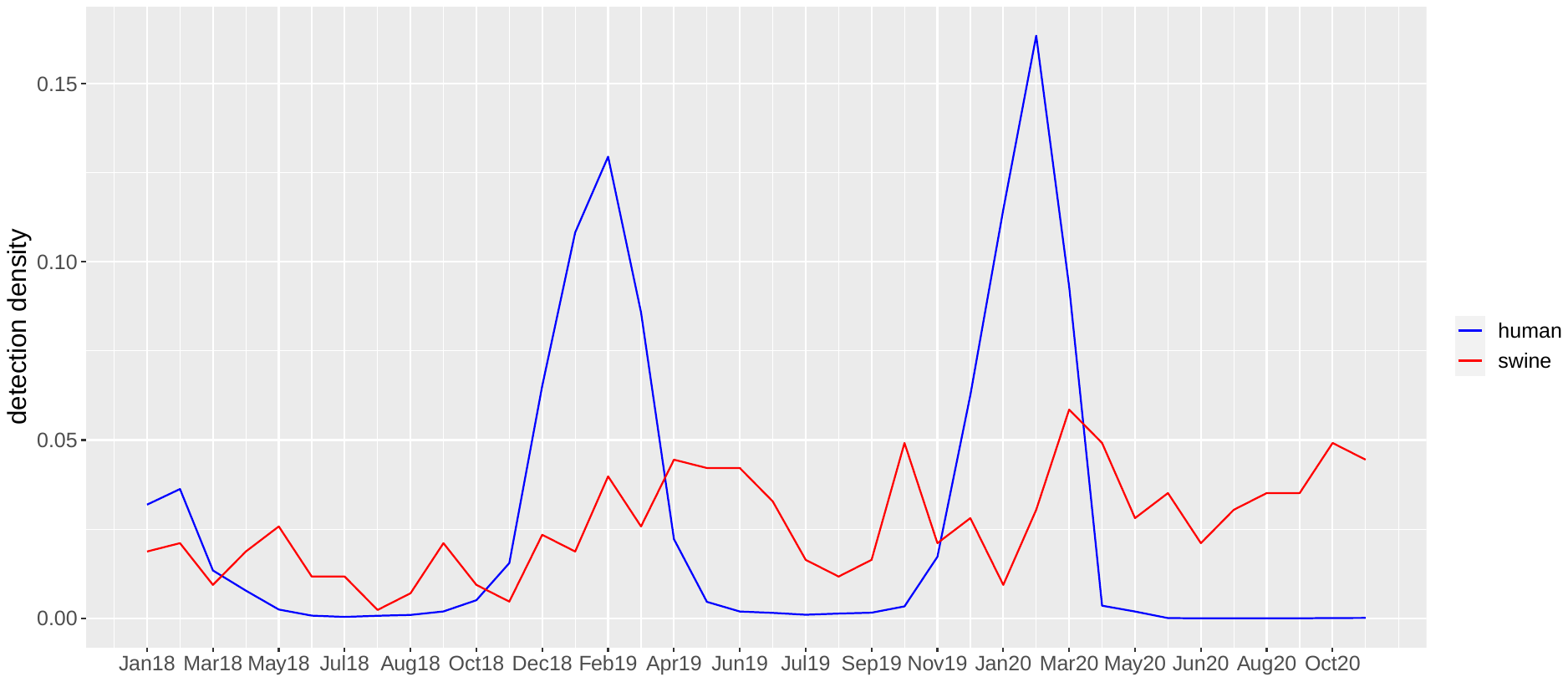

18-19 detection peaks

19-20 detection peaks

Pre-season swine surges

human variant
detection

Figure S1. The normalized pdm09 detection frequency aggregated by 4-week intervals for swine (in red) and humans (in blue) starting January 2018 and ending November 2020. The graph captures two most severe human pdm09 seasons and respective increase in pdm09 circulation in US swine. Arrows show the Fall swine pdm09 detection surges as well as the 18-19 and 19-20 detection peaks in humans and swine. These data capture pdm09 detection frequencies throughout the 18-19, 19-20, and, partially, 17-18 flu seasons. There were consistent September-October swine pdm09 detection surges that occurred prior to the start of the human flu season. These data show that in the 18-19 and 19-20 seasons, the human pdm09 detection peaks preceded the respective swine detection peaks, which corroborates our hypothesis that the swine pdm09 seasons were driven by human-to-swine spillovers. Variant detections during this time period are marked by orange stars.

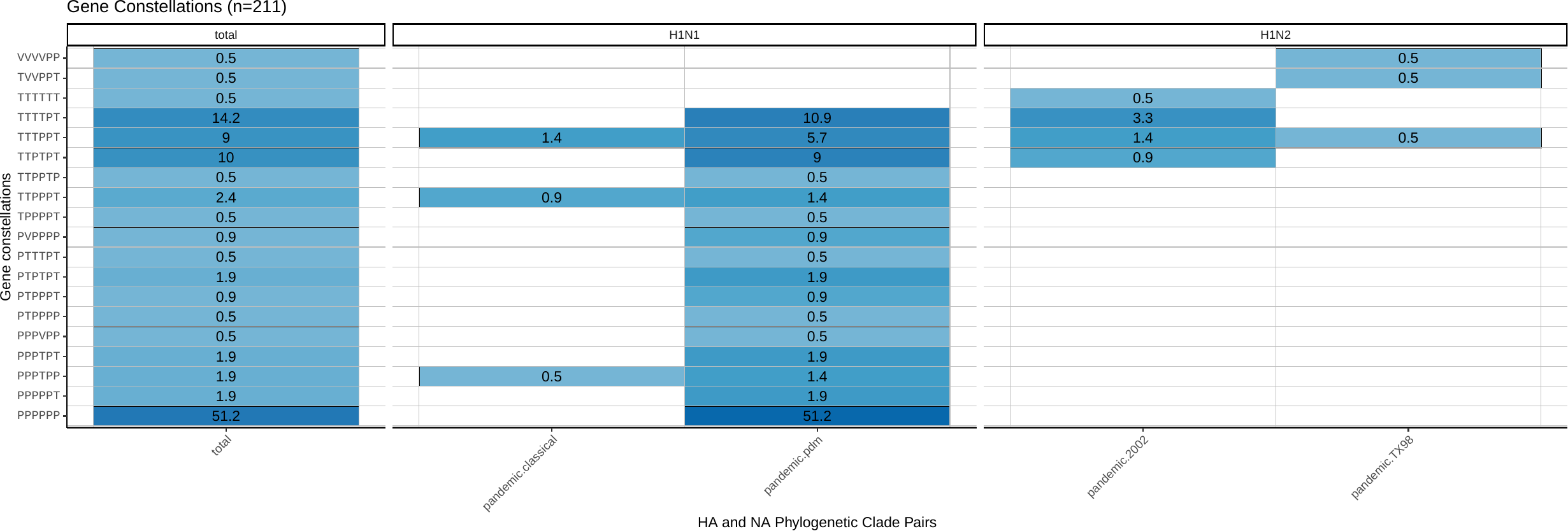

Figure S2. Reassortment patterns of swine H1pdm09 viruses. The data is based on n=211 (2009 to 2021) whole genome sequence IAVs from the USDA influenza A in swine surveillance system. The internal genes are shown in the following order PB2-PB1-PA-NP-M-NS reflecting either the triple-reassortant (T) or H1N1pdm09 (P) evolutionary lineages, or genes derived from the live attenuated influenza vaccine (V)^56^. The 4 most common reassorted internal gene constellations were TTTTPT, TTPTPT, TTTPPT, and TTPPPT, which is consistent with 4 most common internal gene constellation among all H1N1 and H1N2 swine IAVs in the US. The data was pulled and visualized using octoFLUshow^2^.

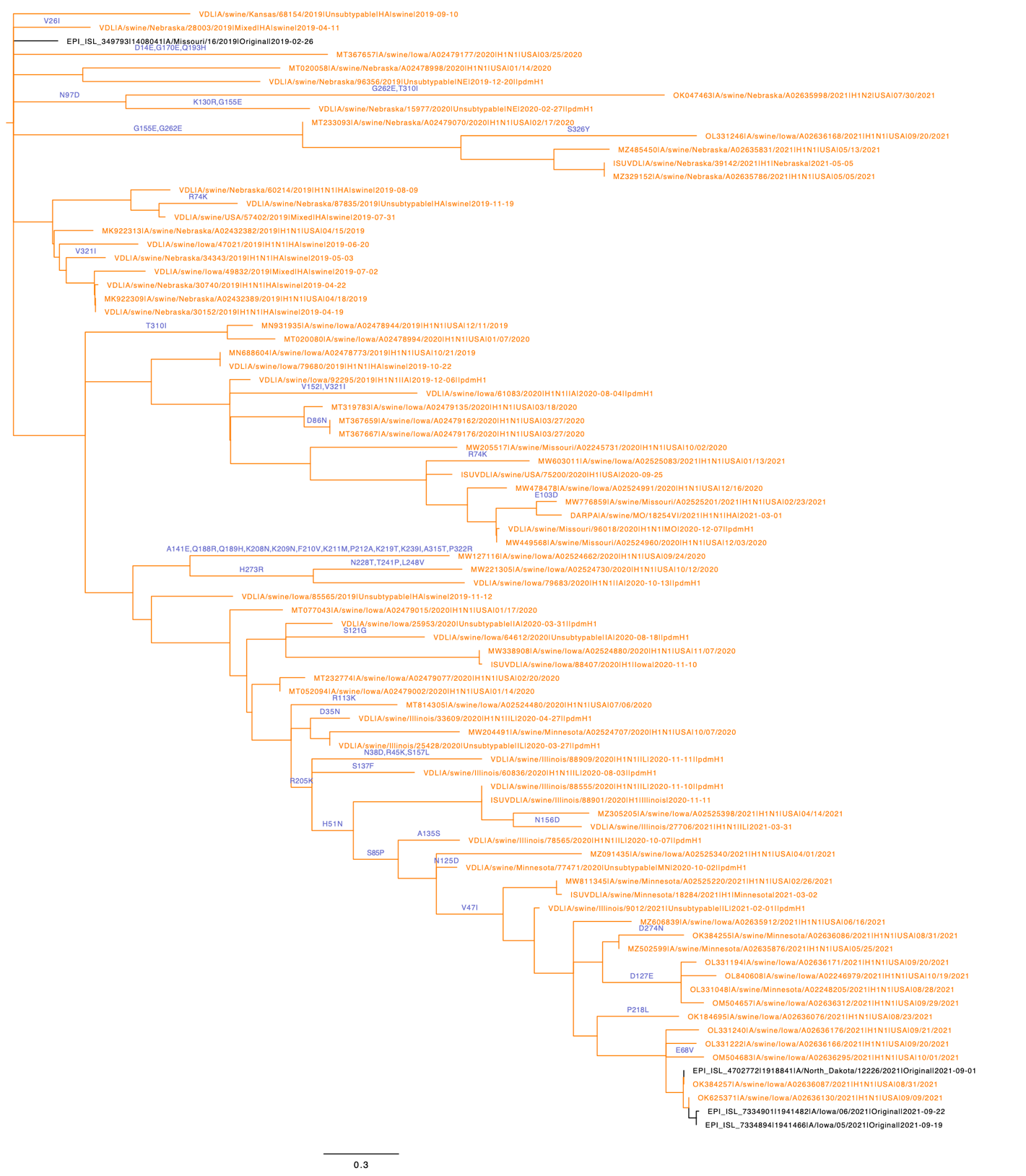

*Figure S3. An HA clade of n=79 swine pdm09 viruses (a 2018-19 human-to-swine spillover) with 4 detections of swine-to-human transmission, 3 of which happened in September 2021 – more than two years after the introduction of the virus into the swine population. The branches are annotated with the respective HA1 amino acid substitutions. Branch-lengths are time-scaled.*

Table S1. Swine H1N1pdm09 strains selected for testing against reference ferret antisera generated against Northern Hemisphere human seasonal H1N1 vaccine strains (A/California/04/2009, A/Michigan/45/2015, A/Brisbane/2/2018, A/Wisconsin/588/2019, and A/Hawaii/70/2019). The strains were selected from different persistent seasonal spillovers by generating an HA1 consensus sequence and then selecting the best-matched field isolate available at the USDA-APHIS Influenza A Virus in Swine virus repository. The internal genes are shown in the following order PB2-PB1-PA-NP-M-NS reflecting either the triple-reassortant (T) or H1N1pdm09 (P) evolutionary lineage.

| Strain name | GenBank Accession | Spillover season | HA1 consensus identity | HA/NA | Internal Gene Constellation |
| --- | --- | --- | --- | --- | --- |
| A/swine/Indiana/A02525081/2021 | MW603033 | 13-14 | 99.7% | H1pdm/N1pdm | PPPPPP |
| A/swine/Colorado/A02635828/2021 | MZ485434 | 15-16 | 100% | H1pdm/N1pdm | TTTTPT |
| A/swine/Kansas/A02248038/2021 | MZ666816 | 17-18 | 99.4% | H1pdm/N1pdm | TTPTPT |
| A/swine/Iowa/A02524480/2020 | MT814305 | 18-19 | 99.7% | H1pdm/N1pdm | TTTPPT |
| A/swine/Indiana/A02635811/2021 | MZ476800 | 19-20 | 99.1% | H1pdm/N1pdm | TTTTPT |
| A/swine/Missouri/A01104146/2020 | MW579260 | 19-20 | 98.5% | H1pdm/N1pdm | PPPPPP |

Table S2. Amino acid difference table over the HA1 subunit for the pdm09 Northern Hemisphere vaccine strains and swine pdm09 representative strains. The amino acid differences are shown relative to A/California/04/2009.

| site | **A/California/04/2009** | **A/Michigan/45/2015** | **A/Brisbane/02/2018** | **A/Wisconsin/588/2019** | **A/Hawaii/70/2019** | A/swine/Indiana/A02525081/2021 | A/swine/Colorado/A02635828/2021 | A/swine/Kansas/A02248038/2021 | A/swine/Iowa/A02524480/2020 | A/swine/Indiana/A02635811/2021 | A/swine/Missouri/A01104146/2020 |
| --- | --- | --- | --- | --- | --- | --- | --- | --- | --- | --- | --- |
| 2 | T |  |  |  |  |  | I |  |  |  |  |
| 45 | R |  | G |  |  |  |  |  |  |  |  |
| 47 | V |  |  |  |  |  | I |  |  |  |  |
| 48 | A |  |  |  |  |  | D |  |  |  |  |
| 68 | E |  |  |  |  | G | G |  |  |  |  |
| 74 | S |  | R | R | R |  |  | R | R | R | R |
| 83 | P | S | S | S | S | S | S | S | S | S | S |
| 84 | S | N | N | N | N |  | N | N | N | N | N |
| 97 | D | N | N | N | N | N | N | N | N | N | N |
| 113 | R |  |  |  |  |  |  |  | K |  |  |
| 120 | T |  |  |  |  |  |  |  |  | A |  |
| 129 | N |  |  | D | D |  | D |  |  | D | D |
| 130 | K |  |  | N |  |  |  |  |  |  |  |
| 137 | P |  |  |  |  |  |  |  | S |  |  |
| 146 | K |  |  |  |  |  | N |  |  |  |  |
| 149 | I |  |  |  |  |  |  | V |  |  |  |
| 152 | V |  |  |  |  |  |  |  |  | I |  |
| 155 | G |  |  |  |  | E |  | E |  |  | E |
| 156 | N |  |  | K |  |  |  |  |  |  |  |
| 161 | L |  |  | I |  |  |  |  |  | I |  |
| 162 | S | N | N | N | N |  | N | N | N | N | N |
| 163 | K | Q | Q | Q | Q | Q | Q |  | Q | Q | Q |
| 164 | S |  | T | T | T |  |  | T | T | T | T |
| 173 | V |  |  |  |  |  |  |  | I |  |  |
| 183 | S |  | P | P | P |  |  | P | P | P | P |
| 185 | S | T | T | I | I | T | T | T | T | I | I |
| 187 | D |  |  |  | A |  |  |  |  |  | S |
| 189 | Q |  |  |  | E |  |  |  |  |  | E |
| 191 | I | L | L | L | L | L | L | L | L | L | L |
| 197 | T | A | A | A | A | A | A | A | A | A | A |
| 203 | S | T | T | T | T | T | T | T | T | T | T |
| 205 | R |  |  |  |  | K | K |  |  |  |  |
| 216 | I | T | T | T | T | K | S | E | T | T | T |
| 222 | D |  |  |  |  |  |  | N |  |  |  |
| 223 | Q | R | R |  |  |  |  |  |  |  |  |
| 239 | K |  |  |  |  | R |  |  |  |  |  |
| 250 | V |  |  | A |  |  |  |  |  | A |  |
| 256 | A | T | T | T | T | T | T | T | T | T | T |
| 260 | N |  |  | D | D |  |  |  |  | D | D |
| 282 | P |  | A |  |  |  |  |  |  |  |  |
| 283 | K | E | E | E | E | E | E | E | E | E | E |
| 295 | I |  | V | V | V |  |  | V | V | V | V |
| 298 | I |  | V |  |  |  |  |  |  |  |  |
| 321 | I | V | V | V | V |  | V | V | V | V | V |
| **Total aadiff** |  | **14** | **21** | **23** | **21** | **14** | **20** | **19** | **20** | **23** | **22** |
